# Cancer Cells in all EMT States Lack Rigidity Sensing Depletion of Different Tumor Suppressors Causes Loss of Rigidity Sensing in Cancer Cells

**DOI:** 10.1101/2022.06.06.495045

**Authors:** Chloe Simpson, Vignesh Sundararajan, Tuan Zea Tan, Ruby Huang, Michael Sheetz

## Abstract

Cancer cells have many different behaviors from epithelial to mesenchymal forms. We report here that 36 distinct tumor cell lines regardless of EMT form or other features lack the ability to sense rigidity and will grow on soft surfaces. In the majority of lines, cells were missing at least one protein needed for rigidity sensing (primarily tropomyosin2.1 (Tpm2.1) but also PTPN12, FilaminA (FLNA), and myosinIIA) while all had high levels of Tpm3. In the few cases where the major rigidity sensing components were present, those tumor cells were not able to sense rigidity. Thus, we suggest that tumor cells can lose the ability to sense rigidity by many different means and that the loss of rigidity sensing is sufficient to cause the transformed phenotype that enables targeted treatments.

## Introduction

For many years, it has been evident that the frequency of cancerous tumors correlates with the frequency of injury/inflammation events (Coussens and Werb, 2002; Il’yasova et al, 2005; Todoric et al, 2016; Michels et al, 2018). In both regeneration and cancer, the blocks to adult cell growth are removed; but in normal regeneration, the blocks are restored (Hart, 2002; Landén et al, 2016). In the case of tumorigenic growth, the blocks to growth remain off and cells have the ability to grow in a variety of environments, even on soft matrices, often with the aid of activating hormones (Feitelson et al, 2015). Early studies of tumor cells showed that they all were able to grow on soft surfaces, i.e. were transformed; whereas normal cells required rigid surfaces to grow (Hamburger and Salmon, 1977; Sheetz et al, 2019). Recent findings support the hypothesis that the removal of the block to growth in cancers and the ability to grow on soft surfaces may be linked. In particular, the transformed state can be induced by depleting rigidity sensors in normal cells from different tissues; and when rigidity sensing is restored to tumor cells, they require rigid surfaces for growth (Gunning et al, 2015; Wolfenson et al, 2016; Qin et al, 2018; Yang et al, 2020). This has led to the hypothesis that all tumor cells are transformed through the loss of rigidity sensing.

In addition, tumor growth is linked to many different signalling pathways and there is a wide range of behaviors of tumor cells that has confounded general treatments of cancers. Often blocking one growth pathway such as caused by mutation of the EGF receptor with an EGFR inhibitor results in the shrinkage of the tumor but metastases often arise with other growth stimuli (Harari, 2004; Nagano et al, 2018). Thus, there is a general ability of tumor cells to grow and they can do so with many different stimuli. This can be understood if tumor cells lack the normal blocks to growth that are removed in wound healing. There are common markers for wound healing and tumor growth in many different systems including an increase in the levels of miR-21 that is highly expressed in brain (Zhang et al, 2018), liver (Marquez et al, 2010), skin (Wang et al, 2012), and even axolotl limb regeneration (Holman et al, 2012). Similarly, many tumor cells have increased miR-21 levels that correlate with the severity of the cancer (Feng and Tsao, 2016; Wu et al, 2017). Since the loss of rigidity sensing enables normal cells to grow, we hypothesize that the loss of rigidity sensors plays a major role in tumor cells and that most if not all tumor cells will lack rigidity sensing.

Evidence from RNA sequencing has shown that the levels of many different mRNAs change upon the loss or restoration of rigidity sensing in cells from diverse tissues (Stefen et al, 2018). Thus, there appears to be a major state change upon transformation or the loss of rigidity sensing that results in general changes in cell behaviour such as a decrease in single cell rigidity (Cross et al, 2007; Swaminathan et al, 2011, Coughlin et al, 2013), increased traction forces (Peschetola et al, 2013; Li et al, 2016, Li et al, 2017), and increased mechanical sensitivity (Betof et al, 2015; Berrueta et al, 2018; Takao et al, 2019; Tijore, et al, 2019). If tumor cells have common properties regardless of the tissue and the exact cause of the tumor, then there may be treatments that would inhibit the growth of a wide diversity of tumors such as mechanical stress.

Alternatively, tumor cells often differ in many respects and one aspect that has been tied with the severity of the disease is the epithelial to mesenchymal transition that often comes with tumor progression in later stage disease. A bank of tumor cells that range from epithelial to mesenchymal has been characterized extensively in terms of morphology and markers for the epithelial and mesenchymal states (Tan et al, 2013; yTan et al, 2014; Tan et al, 2015; Sundararajan et al, 2019). We tested if they all have lost the ability to sense rigidity, which is consistent with our hypothesis that tumor cells should all lack the rigidity sensing whether or not they come from stage 1 or stage 4 tumors. Irrespective of the mutations involved and the phenotype, all the cell lines lacked rigidity sensors and grew on soft surfaces.

Figure 1 illustrates some of the key players linking mechanosensing and cancer. TPM2.1 for example has been reported as silenced and downregulated in breast (Dube et al, 2016), colorectal (Cui et al, 2016) and urothelial bladder cancer (Humayun-Zakaria et al, 2019). In contrast TPM3 is most often overexpressed (Malfatti et al, 2015). FLNA has a complex role in cancer, with both overexpression and loss of expression being associated with poor prognosis (Nakamura et al, 2007; Nomachi et al, 2008; Nakamura et al, 2009; Bedolla et al, 2009; Nallapali et al, 2012; Savoy and Ghosh, 2013; Yue et al, 2013, Nakamura et al, 2014). PTPN12 is a protein tyrosine phosphatase that binds to FLNA by a proline rich domain region, and is known to have interactions with various RTKs including INSR, INSRR, EGFR, as well as GRB2 and cSRC. In triple negative breast cancer restoration of PTPN12 has been shown to suppress EGFR, HER2, and PDGFRβ (Sun et al, 2011), where it acts as a tumour suppressor. In addition, decreased expression has previously been linked to increased motility in Ovarian cancer (Villa-Muruzzi et al, 2011). Finally, myosinIIa has diverse roles in ovarian cancer (Krendel, M. & Mooseker, 2005; Berg et al, 2001; Yoshida et al, 2004; Li and Yang, 2016; Pecci et al, 2018), from increasing contractility in drug resistant tumours cells (Kapoor et al, 2017), clearing mesothelial cells during metastasis (Iwanicki et al, 2011) as well as being associated with cancer cell motility and migration (Vicente-Manzanares et al, 2009; Ouderkirk and Krendel, 2014).

**Figure 1.**
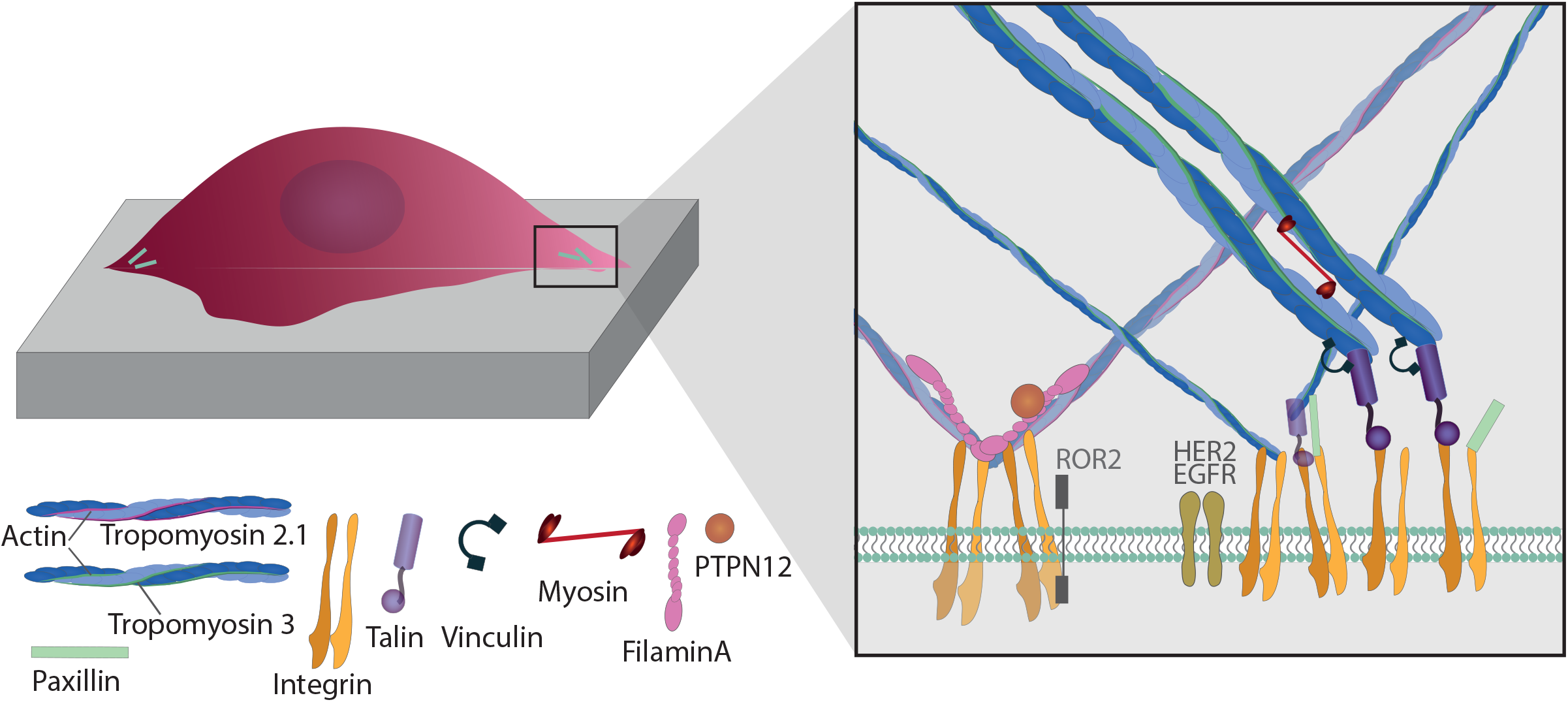
Illustration of mechanosensititve proteins at focal adhesions featuring TPM3, TPM2.1, FLNA, PTPN12 and MyosinIIa. FLNA interacts with integrins and the dimerised FlnA-ig24, actin and the actin binding domain and secondary actin binding sites as well as protein phosphatase PTPN12. MyosinIIa binds at both ends to parallel actin fibres to create regions of acto-myosin contractility. Other well known focal adhesion proteins talin, vinculin and paxillin are also shown.

## Results

We selected a panel of 36 ovarian cancer cells lines of varied origin, one ovarian epithelial cell line and a human foreskin fibroblast cell line. The panel of ovarian cell lines was previously compiled to represent a full spectrum of epithelial to mesenchymal phenotypes (Sundararajan et al, 2019), and originated from various ovarian cancer subtypes, diseases, and stages (Table 1). This panel included 12 cell lines of metastatic origin and 23 cell lines of primary tumour origin, with cells from ovarian serous adenocarcinoma, high grade ovarian serous adenocarcinoma, ovarian endometrioid carcinoma and ovarian cystadenocarcinoma. The panel includes several cell lines from epithelial, intermediate epithelial, intermediate mesenchymal and mesenchymal groups as categorised using methods previously described (Huang et al, 2013; Tan et al, 2014; Tan et al, 2015). We assayed the protein levels of known mechanosensitive proteins TPM2.1, TPM3, FLNA, PTPN12 and Myosin IIa across the panel of cell lines by western blot and quantified these blots (Figure 2A, 2B). Intensity scores from western blots were standardised to a range of 0-1 to show relative expression of each protein, 1 being the highest expression observed across the panel of cell lines. We observed a striking phenotype whereby 18 of the 36 cell lines had no expression of TPM2.1 (Figure 2C), while all had TPM3 at levels equal to or greater than normal cell controls (Figure 2D). A further seven lines, OV90, OV56, TOV112D, HeyA8, Tyknu, SKOV3 and EF021 had no or very low expression of PTPN12 (Figure 2E). One cell line, EF021 had no FLNA (Figure 2F). One cell line, CH1 cells, had no Myosin IIa (Figure 2G). This demonstrates that loss of rigidity sensing proteins occurs in the majority of ovarian cancer cell lines. Co-occurring loss of TPM2.1 and PTPN12 was observed in four cell lines, TYKNu, OVCA420 and TOV112D, SKOV3, and the cell line with no FLNA, EF021 cells, also had low PTPN12. Thus, of the 36 cell lines, the majority had decreased levels of rigidity sensor proteins and all had high levels of Tpm3 that inhibits rigidity sensing (Yang et al, 2020).

**Figure 1.**
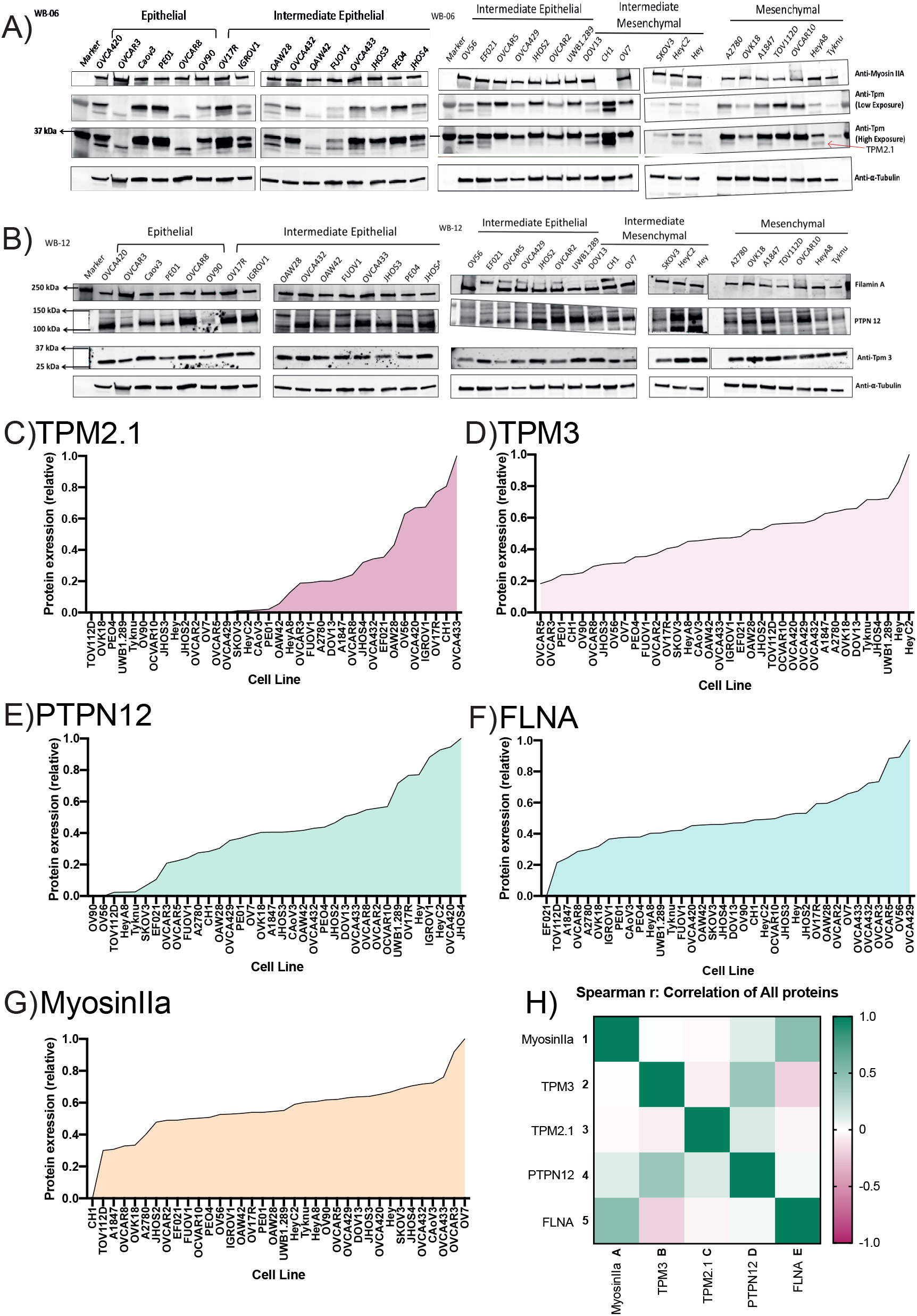
A) Western blots of MyosinIIA and Tropomyosin in order of epithelial to mesenchymal index B) Western blots of FilaminA and TPM3 in order of epithelial to mesnenchymal index. C) Relative, normalised expression level of TPM2.1 in Ovarian cancer cell lines ranked by ascending expression. Eighteen cell lines had no expression TPM2.1. D) Relative, normalised expression level of TPM3 in Ovarian cancer cell lines ranked by ascending expression. All cell lines expressed TPM3. E) Relative, normalised expression level of PTPN12 in Ovarian cancer cell lines ranked by ascending expression. Seven cell lines had very low expression of PTPN12. F) Relative, normalised expression level of FLNA in Ovarian cancer cell lines ranked by ascending expression. One cell line, EF021, did not express FLNA. G) Relative, normalised expression level of MyosinIIa in Ovarian cancer cell lines ranked by ascending expression. One cell line, CH1, had no expression of MyosinIIa. H) Correlation between the expression of all proteins was tested by Spearman rank correlation. PTPN12 and TPM3 were postively correlated, with a R^2^ of 0.414. MyosinIIa and FLNA were positively correlated with and R^2^ of 0.493.

**Table 1.** Panel of ovarian cancer cell lines, annotated with RRID, disease type, tumour stage, and EMT status.

| Cell Line | RRID | Disease | Tumour stage | EMT status |
| --- | --- | --- | --- | --- |
| IOSE523 | CVCL_E234 | Ovarian epithelial (SV40 transformed) | N/a |  |
| BJ1-hTERT | CVCL_6573 | Human foreskin fibroblast | N/a |  |
| CAOV3 | CVCL_0201 | High grade ovarian serous adenocarcinoma | Primary | Epithelial |
| OV90 | CVCL_3768 | Ovarian adenocarcinoma | Metastasis | Epithelial |
| OVCA420 | CVCL_3935 | Ovarian serous adenocarcinoma | Primary | Epithelial |
| OVCAR3 | CVCL_0465 | High grade ovarian serous adenocarcinoma | Metastasis | Epithelial |
| OVCAR8 | CVCL_1629 | High grade ovarian serous adenocarcinoma | Primary | Epithelial |
| PEO1 | CVCL_2686 | Ovarian cystadenocarcinoma | Metastasis | Epithelial |
| EFO21 | CVCL_0029 | Ovarian cystadenocarcinoma | Metastasis | Intermediate epithelial |
| FUOV1 | CVCL_2047 | High grade ovarian serous adenocarcinoma | Primary | Intermediate epithelial |
| IGROV1 | CVCL_1304 | Ovarian endometrioid adenocarcinoma | Primary | Intermediate epithelial |
| JHOS2 | CVCL_4647 | High grade ovarian serous adenocarcinoma | Primary | Intermediate epithelial |
| JHOS3 | CVCL_4648 | Ovarian serous adenocarcinoma | Primary | Intermediate epithelial |
| JHOS4 | CVCL_4649 | High grade ovarian serous adenocarcinoma | Primary | Intermediate epithelial |
| OAW28 | CVCL_1614 | High grade ovarian serous adenocarcinoma | Metastasis | Intermediate epithelial |
| OAW42 | CVCL_1615 | Ovarian cystadenocarcinoma | Metastasis | Intermediate epithelial |
| OV17R | CVCL_2672 | Ovarian adenocarcinoma | Primary | Intermediate epithelial |
| OV56 | CVCL_2673 | Ovarian serous adenocarcinoma | Metastasis | Intermediate epithelial |
| OVCA429 | CVCL_3936 | Ovarian cystadenocarcinoma | Primary | Intermediate epithelial |
| OVCA432 | CVCL_3769 | Ovarian serous adenocarcinoma | Primary | Intermediate epithelial |
| OVCA433 | CVCL_0475 | Ovarian serous adenocarcinoma | Primary | Intermediate epithelial |
| OVCAR2 | CVCL_3941 | Ovarian carcinoma | Metastasis | Intermediate epithelial |
| OVCAR5 | CVCL_1628 | Ovarian serous adenocarcinoma | Primary | Intermediate epithelial |
| PEO4 | CVCL_2690 | Ovarian cystadenocarcinoma | Metastasis | Intermediate epithelial |
| UWB1.289 | CVCL_B079 | Ovarian carcinoma | Primary | Intermediate epithelial |
| A2780 | CVCL_0134 | Ovarian endometrioid adenocarcinoma | Primary | Intermediate mesenchymal |
| CH1 | CVCL_4992 | Ovarian mixed germ cell tumor | Metastasis | Intermediate mesenchymal |
| DOV13 | CVCL_6774 | Ovarian adenocarcinoma | Primary | Intermediate mesenchymal |
| HEY | CVCL_0297 | High grade ovarian serous adenocarcinoma | Primary | Intermediate mesenchymal |
| HEYC2 | CVCL_X009 | High grade ovarian serous adenocarcinoma | Primary | Intermediate mesenchymal |
| OV7 | CVCL_2675 | Ovarian carcinoma | Primary | Intermediate mesenchymal |
| SKOV3 | CVCL_0532 | Ovarian serous cystadenocarcinoma | Metastasis | Intermediate mesenchymal |
| HEYA8 | CVCL_8878 | High grade ovarian serous adenocarcinoma | Primary | Mesenchymal |
| OVCAR10 | CVCL_4377 | Ovarian carcinoma | Primary | Mesenchymal |
| OVK18 | CVCL_3770 | Ovarian endometrioid adenocarcinoma | Metastasis | Mesenchymal |
| TOV112D | CVCL_3612 | Ovarian endometrioid adenocarcinoma | Primary | Mesenchymal |
| TYKNU | CVCL_1776 | High grade ovarian serous adenocarcinoma | Primary | Mesenchymal |

Overall, no correlation was shown across the panel with epithelial to mesenchymal status and the expression of any of the assayed proteins across patient samples from CSIOVDB (Tan et al, 2015; Supplementary Figure 1A-E) or in this panel of cell lines (Supplementary Figure 2A), indicating that the loss rigidity sensing is an early event in metastatic progression, occurring before EMT. A significant correlation was observed between expression of FLNA and MyosinIIA (Spearman r 0.4493, P = 0.002) and between PTPN12 and TPM3 (Spearman r 0.414, p = 0.012) (Figure 2G, Supplementary Figure 2B). We additionally assayed a few other known mechanosensitive proteins in 33 of the cell lines, TPM1, TPM2 (including TPM2.1 and other splice variants), DAPK, Piezo1 and mir21 RNA. All of the cell lines expressed these proteins (Supplementary Figure 3A-E) and additional correlation was observed between DAPK and TPM1 expression (Spearman r 0.460, p = 0.007) (Supplementary Figure 3F).

We selected 3 tumor lines from the panel which retained all the proteins assayed (Figure 3A) and tested whether they had the ability to detect changes in substrate stiffness phenotypically. The three cell lines were IGROV1 which is an ovarian endometriod carcinoma cell line, OVCA433, and OVCA433, which are both ovarian serous adenocarcinoma cell lines, IOSE523 and OVCAR3 cell lines (Figure 3A). We also included two negative control lines, a human foreskin fibroblast cell line known to be capable of effective rigidity sensing BJ-3ta (Yang et al, 2019) and an ovarian epithelial cell line transformed using SV-40, IOSE523. We additionally included a positive control, a high grade ovarian serous carcinoma cell line which lacks TPM2.1, OVCAR3. As cells spread on the surface they tested the stiffness by contracting matrix adhesions (Byron et al, 2010; Hu et al, 2017). This sarcomeric pinching was quantified by plating cells on PDMS micropillars coated with fibronectin and analyzing pillar displacements over time (Ghassemi et al, 2012). As cells spread, the pillars were deflected in response to forces applied by the cell. Rigidity-sensing events were defined as contractile deflections of two pillars simultaneously by >30 nm for >20s such as those of pillar 1 and 2 (Figure 3Biii) whereas in many cases pillar deflections were not correlated (pillars 1 and 2 in Fig. 3Biv). The number of events per unit area per unit time was a measure of the cell’s ability to sense rigidity. In this case, cells were plated on 0.8 μm pillars and imaged for 30 minutes, at a frame rate of 1 per second. An automated program determined the number of contractile events that fit the criteria for rigidity-sensing events from movies of the pillars covered by cells (see rigidity-sensing pillar deflections shown in green and non-correlated pillar deflections shown in red (Figure 3i).

**Figure 3.**
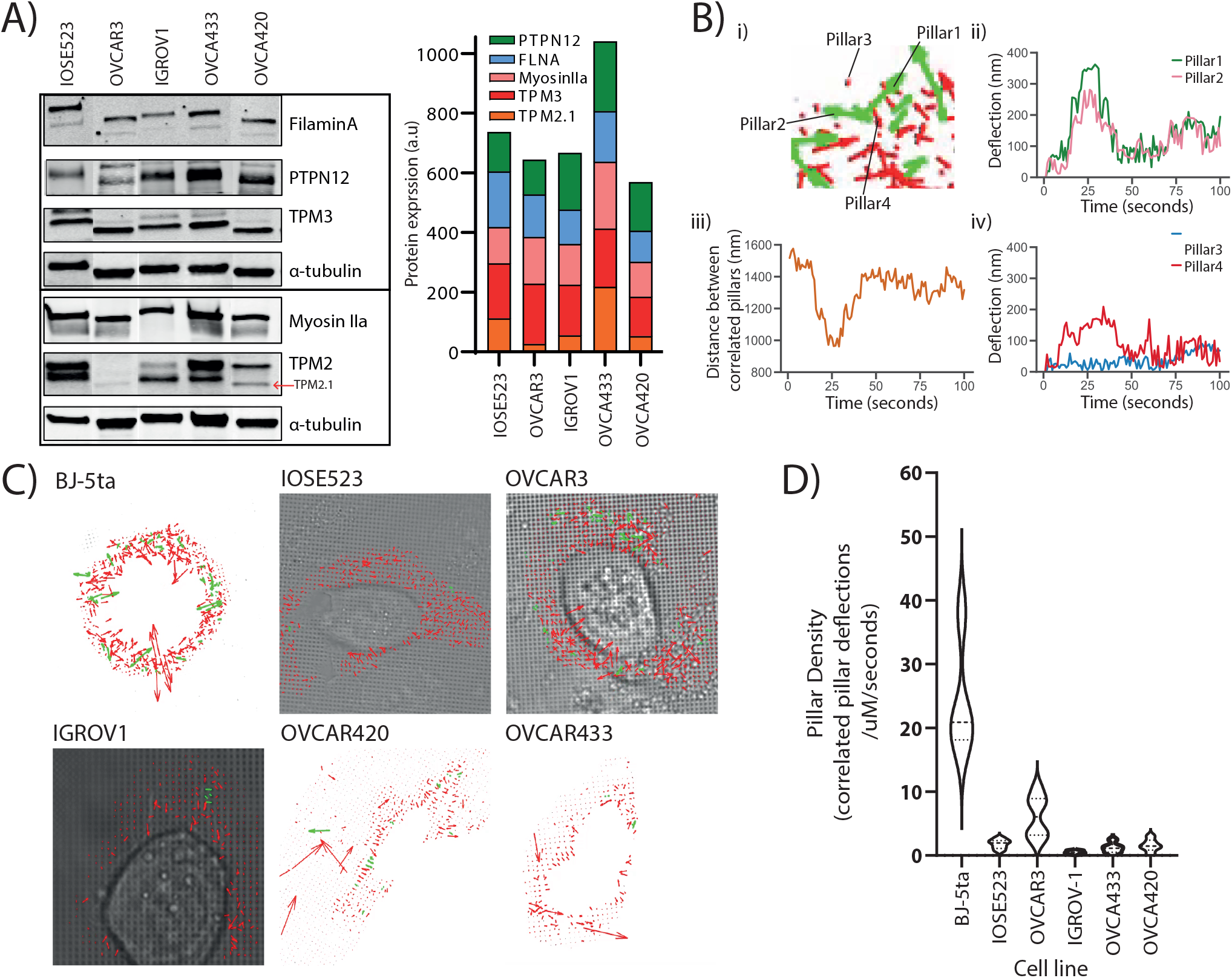
A) Western blots of selected cell lines showing FilaminA, PTPN12, TPM3, and a-tubulin (normalisation control). Western blot of selected cell lines showing MyosinIIa, TPM2 and a-tubulin (normalisation control). TPM2.1 isoform is indicated with a red arrow. OVCAR3 had no expression of TPM2.1. Cell lines were selected that retained all assayed proteins, TPM3., TPM2.1, FLNA, PTPN12 and MyosinIIa, along with sswith ovarian epithelial cell line IOSE523 and OVCAR3, which lacks TPM 2.1 B) i) Close up image of deflections of four pillars, taken from still from BJ cells. ii) Pillar 1 and 2 are correleted iii) Distance between pillars 1 and 2 iv) pillar 3 and 4 were not correlated. C) Stills from movies generated through the pilartracker plug in for ImageJ from MBI and matlab script to analyse correlated pillar deflections, so termed contractile pairs. Cells were plated on 0.8um micropilars and imaged at 1 frame per second for 30 minutes. D) Pillar density shown for ech cell line. Pillar density is calculated by number of pillars per area (uM^2^) per time (seconds) each cell line. Significant difference between BJ cells and all other cell lines, determined by one way ANOVA with Tukey correction.

The movies from this imaging were processed by removing drift using the ImageJ plugin stackreg with translation, detecting and tracking pillars with the ImageJ plugin Pilartracker and calculating the correlated pillar deflections using a custom script for Matlab. Based upon the unbiased computer analysis, the HFF (BJ-5ta) cells had a density of rigidity sensing 24.27 events per second per μm^2^ (N=4, Figure 3D). In contrast, all the ovarian cells lines had a significantly lower number of contractile events. IOSE523 cells had a mean of 1.8 (n=7) contractile pairs per second per μm^2^, OVCAR3 cells had a mean of 6.13 (n=6), IGROV1 cells had a mean of 0.49 (n=5), OVCA433 cells had a mean of 1.18 (n=5) and OVCA420 (n=5) cells had a mean of 1.57 contractile units per second per μm^2^, where in is the number of individual cells. Thus, all of the tumor cells, and the transformed ovarian epithelial cell line lacked rigidity sensing contractions, which further supported the hypothesis that tumor formation involved the loss of rigidity sensing.

Previously it has been demonstrated that one marker of cells capacity to detect changes in substrate stiffness is reflected in how their actin is aligned (Prager-Khoutorsky et al, 2011; Gupta et al, 2015, Gupta et al, 2016; Sarangi et al, 2017). Non-transformed cells on soft surfaces have a low alignment of actin and do not form polarised stress fibres, while on rigid substrates actin fibres align in a coordinated direction and the cell polarises. Using this criterion, we tested the alignment of actin filaments by imaging f-actin using a phalloidin stain and calculating the nematic order parameter of each cell on two PDMS substrates of different stiffness, 2MPa and 5kPa (Prager-Khoutorsky et al, 2011). To quantify actin ordering, images were cropped to single cell images and nematic order parameter was measured using a custom Matlab script (Gupta et al, 2019). This gave a single numerical value for the alignment score of the actin filaments between 1 and 0, 1 being all fibres in the same direction and 0 being no fibres in the same direction. In non-transformed fibroblasts, a significant difference in actin alignment was observed between 5kPa and 2MPa. On stiffer substrates, fibroblasts had a more ordered actin structure with higher polarization but on softer substrates, they had a more disordered actin (Figure 4A, Gupta et al, 2019). Significantly higher nematic order parameter was observed in the negative control BJ-3ta HFF cells plated on 2MPa compared to 5kPa. In contrast, none of the ovarian cell lines demonstrated a significant difference in the nematic order of actin between soft and rigid substrates (Figure 4B/C), including the ovarian epithelial line IOSE523. This ovarian epithelial line, although not cancerous in origin, is SV-40 transformed and has lost the capacity to sense substrate stiffness (Chang et al, 2013).

**Figure 4.**
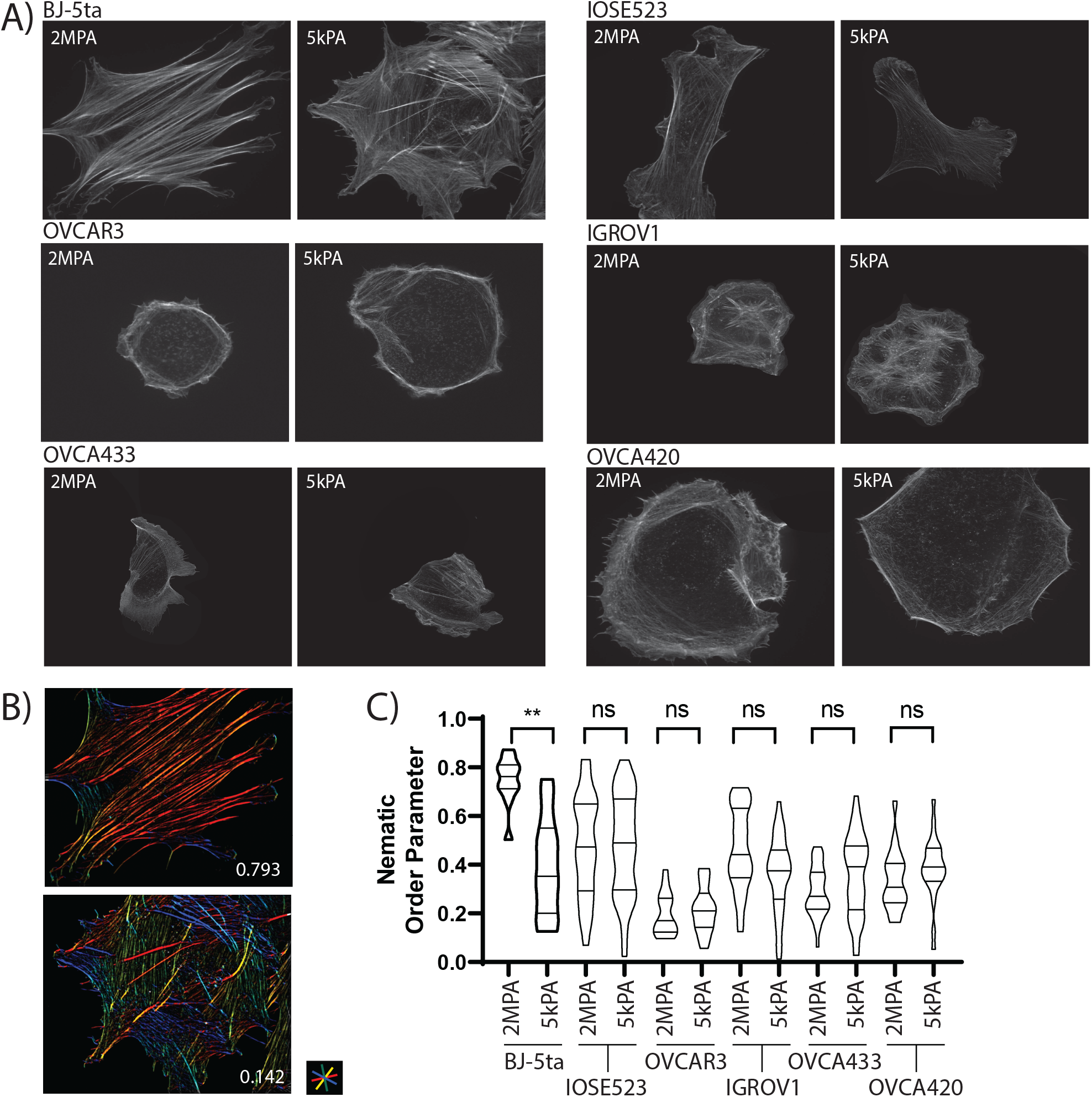
Nematic order perameter analysis demonstrates that none of the ovarian cell lines display rigidity sensing between soft and rigid surfaces. A) Images of Phalloidin staining for f-actin in six cell lines on PDMS surfaces of 2MPa and 5kPa stiffness. B) Example images of actin alignment quantification in HFF (BJ-5ta) cells on 2MPa (top) and 5kPa (bottom) with the nematic order perameters of 0.793 and 0.142 respectively. Images are output from the custom matlab script and are colour coded by actin fibre angle C) Quantification of the nematic order perameter in each cell line on both surfaces. There was a significantly higher nematic order peramter on 2MPa PDMS in HFF cells vs HFF cells on 5kPa PDMS. No other cell line had significant differences in nematic order peramter between 2MPa and 5kPa PDMS.

Through these two methods, measuring the correlated pillar deflections in spreading cells and the nematic order perameter of actin after cell spreading, we show that the remaining cell lines that retained all the proteins assayed phenotypically were unable to sense a change in substrate stiffness in actin alignment, nor were they able to pinch micropillars during the spreading phase of attachment. These cell lines we propose have lost rigidity sensing by currently unknown means.

## Discussion

In this study, all of the ovarian tumor lines tested lack the ability to sense rigidity. There are many differences in the levels of expression of mechanosensory proteins but about 75% of the lines are missing one of the proteins known to be needed for rigidity sensing. Half of the cell lines lack TPM2.1, 20% are depleted of PTPN12 and there are occasional losses of FilaminA or Myosin IIA; but in several cases the basis of the loss in rigidity sensing is unknown. However, the fact that all lines appear to lack rigidity sensing is consistent with the hypothesis that transformation is necessary for tumor formation and transformation results from the loss of rigidity sensing.

Because the rigidity sensor is a complex modular machine with many different components, it is expected that the loss of any one of many different components could block rigidity sensing. Studies have found that rigidity sensing is lost and transformed growth is gained upon the depletion of at least eight different proteins (Lin et al, 2015; Meacci et al, 2016; Maslikowski et al, 2017; Saxena et al, 2017; Wolfenson et al, 2018; Yang et al, 2019). Like any complex process that involves many proteins, there are many possible ways to block function. This is consistent with the general diversity of mutations that give rise to cancer. What is not considered here are the many developmental changes that are involved in large tumor growth and metastasis. Progression in cancer is enabled by transformation, since with some growth, cells can evolve or develop the best systems for growth in their microenvironment. Thus, although the loss of function in any one of a large group of proteins can cause transformation, an even larger number of alterations are needed for tumor growth or metastasis.

TPM2.1 has already been shown to be downregulated in breast (Dube et al, 2016), colorectal (Cui et al, 2016) and urothelial bladder cancer (Humayun-Zakaria et al, 2019). Here we show that TMP2.1 is commonly downregulated or lost in ovarian cancer cells, further providing evidence for TMP2.1 as a tumour suppressor. TPM3, however is most often overexpressed, and is rarely lost throughout this panel of ovarian cancer cell lines, providing evidence of a potential role as an oncogene. From the many different phenotypes of cancer cells even with the loss of rigidity sensing proteins, the common mechanosensing defects in cancer cells are not evident. In particular, the epithelial to mesenchymal transition may increase cell motility and the risk of metastasis or severe disease, however, the loss of rigidity sensing or transformation appears to be a common feature of nearly all tumor cells irrespective of EMT.

The transformed phenotype involves more than the ability to grow on soft agar. Studies of tumor cells, indicate that in general they are softer (more easily deformed), they generate higher traction forces and they are damaged by mechanical perturbations. This is likely only a partial list of the complex functions that are altered with transformation since the mRNA levels of several hundred proteins are altered by the simple expression of TPM2.1 in tumor cells or the depletion of TPM2.1 in fibroblasts. Transformation appears to be a state change that involves a concerted alteration of a characteristic set of functions.

Becoming insensitive to substrate stiffness has several advantages for a cancer cell, firstly they are able to ignore cues in their environment which normally signal them to slow down proliferation, or activate them for apoptosis. It has previously been shown that, in noncancer cells, plating on soft surfaces initiates a signalling cascade for caspase mediated apoptosis (Bosch et al, 2011; McIlwain et al, 2013; Qin et al, 2018) in losing mechanosensitive proteins this signal is ablated or at least weakened meaning that cancer cells can avoid death by avoiding this signalling pathway. Secondly, as cancer cells metastasize, it is conceivable that they will encounter various substrates from collagen to vascular systems and others (Morgan-Parkes, 1995; Stoletov et al, 2010; Chiang et al, 2016; Zavyalova et al, 2019), and being able to adapt to changing mechanical environments would make them more likely to survive to produce metastatic tumours.

The general changes in functions that are common to transformed cells and most if not all tumor cells are potentially very important in indicating ways to inhibit growth of many different tumor cells. In this regard, the mechanical sensitivity of transformed cells is potentially interesting. Not only the fluid shear of tumor cells but also mechanical stretching causes apoptosis. Further, there are indications that mechanical perturbation of tumors in vivo will also inhibit their growth. Other aspects of the transformed state have not been systematically analysed to determine if other treatments would inhibit growth of tumor cells but not their normal cell neighbors.

In summary, these findings show that the transformed state is a common characteristic of a diverse set of tumor cells that show many different behaviors and that the transformed state correlates with the loss of rigidity sensing. This supports the hypothesis that transformation is necessary but not sufficient for tumor growth. Further, the characteristic changes in functions upon transformation may be exploited to aid in the selective inhibition of transformed cell growth.

## Methods and Materials

### Cell culture

BJ-5ta, IOSE523, OVCA433 and OVCA420 cells were cultured in DMEM with high glucose and supplemented with 10% FBS. OVCAR3 and IGROV1 cell lines were cultured in RPMI media supplemented with 10% FBS. Cells were incubated at 37°C, with 80% humidity and 5% C0_2_.

### Micropillar preparation and imaging

Micropillar production was done using precut silicone moulds. PDMS and crosslinker were mixed thoroughly in a ratio of 10:1 and degassed for 30 minutes at 10MB. PDMS mixture was poured onto the mould degassed for 10 minutes at 10MB. The mould was upturned onto a plasma cleaned glass bottomed plate, with a weight on top and further degassed for 10 minutes. The plate-mould-PDMS was then cured for 3 hours at 80°C. The PDMS was demoulded in isopropanol and washed 5x with PBS, or until no isopropanol remained. Before plating the PDMS micropillars were incubated with fibronectin at 37°*C* for 1 hour. PDMS was replaced with cell culture media and cells were plated, and incubated for 30 minutes before imaging. Cells were imaged in the DIC channel using EZ-live Olympus widefield microscope for 30 minutes at 1 frame per second.

TIFF movies of the cells spreading on micropillars were corrected for imaging drift using the FIJI plugin stackreg with translation. Pillars were detected and deflections calculated using the Fiji plugin Pilartracker from MBI. Correlated pair deflections were assigned using the custom matlab script.

### PDMS surface preparation and imaging

To measure nematic order perameter of the actin fibres of cells, PDMS surfaces were produced using Sylgard 184 elastomer kit with varying ratios of elastomer to curing agent. For 2MPa surfaces, the elastomer to curing agent ratio was 10:1, for 5kPa surfaces the ratio was 75:1. Elastomer mixes were spin-coated onto plasma cleaned glass coverslips using a protocol of 200RPM for 10 seconds and 1000RPM for 1 minute. Slides were cured at 80°C for 3 hours. Slides were incubated with fibronectin for 1 hour at 37°C before cells were plated.

### Fluorescence microscopy and analysis

Cells were trypsinised and plated onto both 2MPa and 5kPa slides and incubated for 6 hours at 37°C, at which point they were fixed in 4% Paraformaldehyde for 12 minutes at 37°C. Cells were permeablised with 0.5% Triton in PBS for 10 minutes at room temperature (RT) and blocked with 2% bovine serum albumin for 2 hours at RT. Primary antibody for paxillin (ab32084, abcam) was diluted in 2% BSA at a ratio of 1:1000 and incubated at RT for 2 hours. Secondary antibody Alexafluor-647 anti rabbit at a dilution of 1:1000 and Phalloidin stain (A12379, Thermofisher) for f-actin at a dilution of 1:400 were mixed in 2% BSA added to slides and incubated for 1 hour at RT. The slides were washed 3x in PBS between each step. Cover slips were mounted onto glass slides using Dako mounting medium. Slides were imaged on W1 Live-SR spinning disk super resolution, and deconvolved to super resolution using the inbuilt LIVE-SR feature.

### Western blot analysis

Whole cell protein lysates were used from the previously described ovarian cancer cell line library (SGOCL) (Sundararajan et al., 2019). Cell lysates harvested in RIPA buffer were resolved through SDS-PAGE, followed by blotting on PVDF membranes. Immunoblots were subsequently incubated with appropriate primary antibodies: anti-Filamin-A (ab111620, Abcam); anti-Myosin-IIA (M8064, Sigma); anti-Tropomyosin 2 (T2780, Sigma); anti-Tropomyosin 3 (ab180813, Abcam), anti-α-Tubulin (ab7291, Abcam) diluted in 2% BSA in TBST. Secondary antibodies from Li-COR Biosciences were used: IRDye 800CW goat anti-mouse/rabbit (926-32210, 926-32211), IRDye 680LT goat anti-mouse/rabbit (926-68020, 926-68021). Immunoblots were scanned using the Odyssey Infrared Imaging System (Li-COR) and were converted to gray scale.

