## Supplementary Figure 1. for "Cancer Cells in all EMT States Lack Rigidity Sensing Depletion of Different Tumor Suppressors Causes Loss of Rigidity Sensing in Cancer Cells"

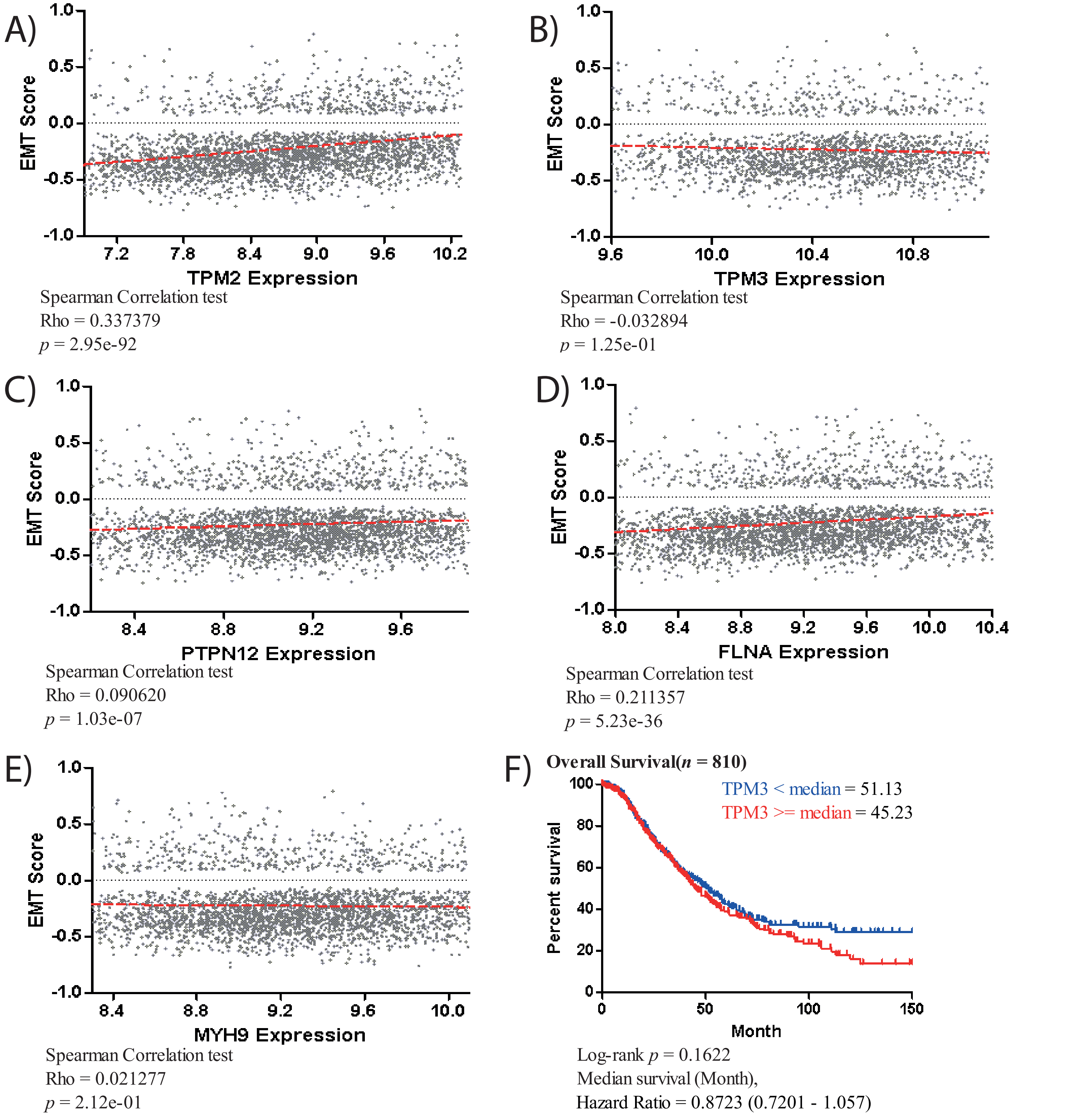


Supplementary Figure 1 A) Correlations of TPM2 with EMT score taken from CSIOVDB, a database of expression levels in clinical samples. B) Correlation of TPM3 with EMT score taken from CSIOVDB C) Correlation of PTPN12 with EMT score taken from CSIOVDB. D) Correlation of FLNA with EMT score taken from CSIOVDB. E) Correlation of MYH9 with EMT score taken from CSIOVDB. None of the proteins had a correlation > 0.35 or <-0.35 for expression vs EMT score. F) Kaplan Meier graph for ovarian cancer patients in CSIOVDB segregated by TPM3 expression. Patients with high TPM3 expression had worse outcome that those with low TPM expression with a hazard ratio of 0.8723.
