## Supplementary Figure 2. for "Cancer Cells in all EMT States Lack Rigidity Sensing Depletion of Different Tumor Suppressors Causes Loss of Rigidity Sensing in Cancer Cells"

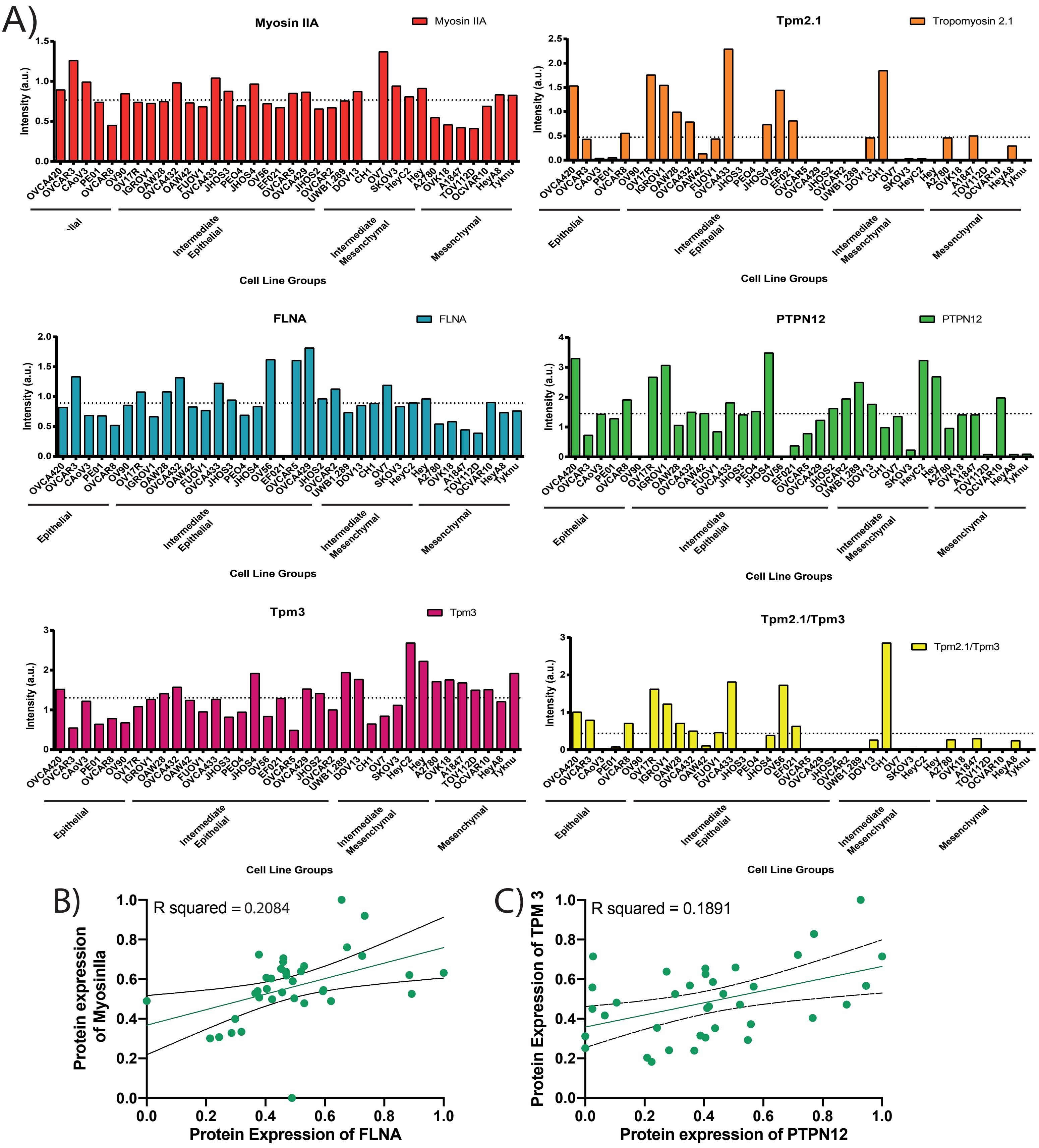


Supplementary Figure 2A) Quantification of each protein, ranked by epithelial to mesenchymal index, also showing the ratio of TPM2.1/TPM3 ranked in the same order. B) Linear regression model for correlation between FLNA and MyosinIIa, the line for the linear regression model has the equation Y = 0.3060*X + 0.3584, and an R squared of squared of 0.2084. Linear regression model for correlation between PTPN12 and TPM3, the line has the equation Y = 0.3914*X + 0.3680, and an R squared of 0.1891
