## Supplementary Figure 3. for "Cancer Cells in all EMT States Lack Rigidity Sensing Depletion of Different Tumor Suppressors Causes Loss of Rigidity Sensing in Cancer Cells"

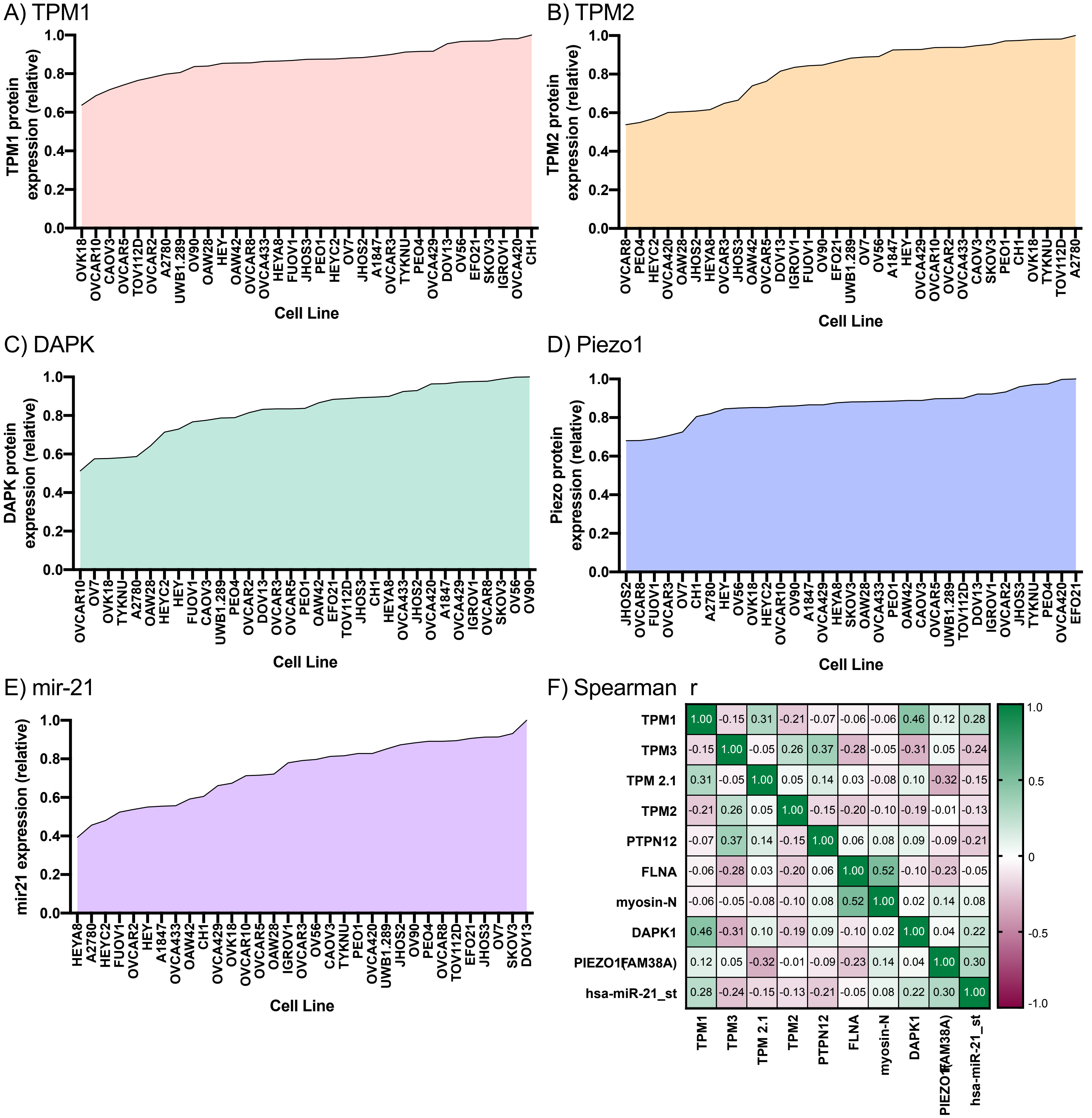


Supplementary Figure 3. Additional proteins and RNA assayed for 33 Ovarian cancer cell lines. Expression levels are relatiewe, within a scale of 0 -1 A) Relative TPM1 protein level in ascending order B) Relative TPM2 protein level in ascending order C) Relative DAPK protein level in ascending order D) Relative Piezo1 protein level in ascending order E) Relative mir21 RNA expression level in ascending order. F) Correlations between expression levels of all assayed proteins in 33 cell lines, determined by Spearman rank correlation. Significant correlation was observed between DAPK and PTPN12 (p = 0.007), PTPN12 and TPM3 (p = 0.036) and myosin-N and FLNA (p =0.002).
